# Same but different: New insights on the correspondence between subjective affective experience and physiological responses from representational similarity analyses

**DOI:** 10.1101/2021.04.30.442153

**Authors:** C. Ventura-Bort, J. Wendt, M. Weymar

**Affiliations:** Department of Biological Psychology and Affective Science, Faculty of Human Sciences, University of Potsdam, 14476 Potsdam, Germany; Faculty of Health Sciences Brandenburg, University of Potsdam, 14476 Potsdam, Germany

## Abstract

Classical views suggest that experienced affect is related to a specific bodily response (*Fingerprint hypothesis*), whereas recent perspectives challenge this view postulating that similar affective experiences rather evoke different physiological responses. To further advance this debate in the field, we used representational similarity analysis (N= 64) to investigate the correspondence between subjective affect (arousal and valence ratings) and physiological reactions (skin conductance response [SCR], startle blink response, heart rate and corrugator activity) across various emotion induction contexts (picture viewing task, sound listening task and imagery task). Significant similarities were exclusively observed between SCR and arousal in the picture viewing task. However, none of the other physiological measures showed a significant relation with valence and arousal ratings in any of the tasks. These findings tend to support the *populations hypothesis*, suggesting that there is no clear match between the evoked physiological responses and the experienced subjective affect between individuals.

**Statement of relevance:** The subjective affective experience evoked by an event is accompanied by physiological responses. The correspondence between physiological response patterns and the experienced affect, however, is still under debate. Classical views (*Fingerprint hypothesis*) suggest that affect is related to a specific physiological response, whereas recent perspectives (*Populations hypothesis*) challenge this view, postulating rather different physiological responses. In the current study, we used representational similarity analysis (RSA) to examine the relation between affective experience, assessed using valence and arousal ratings, and the evoked physiological reactivity across three affect-inducing contexts. Results showed significant similarities exclusively between SCR and arousal in the passive picture viewing task. However, none of the other physiological measures showed a significant relation with valence and arousal ratings in any of the tasks, supporting the populations hypothesis. These findings invite to reframe the relation between physiology and affect from invariant and homogeneous to variant and context-dependent.

## Introduction

> “*Abstract the bodily symptoms from any strong emotion and it goes with them*.”
>
> — William James (1890)

It is long known that strong subjective affective feelings evoked by a particular event are accompanied by prominent physiological reactions (e.g., heart pounding, hand sweating). Bodily changes during ongoing experiences are an important aspect when determining that an event is pleasant, a sound feels activating, or a memory is terrifying. Despite this co-occurrence, the extent of correspondence between physiological reactivity and the experienced subjective affect is still unclear. Theoretical proposals of affect converge on two basic characteristics of an affective subjective experience, i.e., valence, referring to its hedonic value (ranging from pleasant to unpleasant), and arousal, referring to degree of activation (ranging from feeling calm to active, excited) (e.g. Barrett & Russell, 1999; Cacioppo & Berntson, 1994; Kuppens et al., 2013; Lang et al., 1990; Reisenzein, 1994). However, these views diverge in terms of the proposed relation between physiological changes and the valence and arousal properties of affect.

More classical views (also known as the *Fingerprint hypothesis*; Hoemann et al., 2020; Siegel et al., 2018) state that the experienced affect is the by-product of the activation of phylogenetically evolved brain circuits whose function is to ensure the survival of the individual (Barrett, 2017; Cacioppo & Berntson, 1994; Friedman, 2010; Lang et al., 1993; Witvliet & Vrana, 1995). These circuits also orchestrate specific physiological changes that promote adaptation by facilitating the detection and preparation of the organism for overt actions in the face of life-sustaining or life-threatening events (e.g. Lang & Bradley, 2010). From these perspectives, a strong correspondence should exist between the experienced affect and the evoked physiological response, given their overlapping origin. Although the characteristics of the context in which the relevant event is may determine the pattern of physiological changes that ensure the best adaptive response (i.e., fighting vs. flying), if maintained constant, different events evoking similar affective experiences are expected to produce similar physiological responses within and across individuals.

More recent views (also named the *Populations hypothesis*; (Barrett, 2017; Hoemann et al., 2020) suggest that the basic properties of valence and arousal are created based on an assembly of heterogeneous context-dependent instances that vary from each other in their physical characteristics. In this regard, previous studies have shown that the mental representation of affect may be based not so much on the specific characteristics of a particular stimulus, but on similarities within more abstract representations of the properties of affect such as their hedonic value (Chikazoe et al., 2014; Jin et al., 2015). According to this view, due to the physical heterogeneity across events, the physiological demands in each of them will vary in a situated fashion, independently of the associated affective experience. Empirical support comes from a recent experience sampling study by Hoemann and colleagues (Hoemann et al., 2020; see also, Siegel et al., 2018) who investigated whether specific physiological patterns exist for the reported valence and arousal within and across participants. In this study, participants’ subjective affective experience of physiologically triggered events were assessed during a 2-week period. Results revealed variability rather than specificity in the relation between valence and arousal ratings and the detected physiological patterns. That is, a particular physiological pattern was labeled with terms assigned to different levels of valence (e.g. happy, bored) and arousal (e.g. excited, calm), and the same terms were related to two or more physiological patterns (Hoemann et al., 2020). These findings support the *populations* view that the properties of affect are abstract categories that are context-specific, and therefore unrelated to the physical demands of a particular situation (Hoemann et al., 2020). However, due to the ecological adaptation of the design, the environment in which the samples were collected may have considerably varied from participant to participant. Furthermore, the used physiological variables mostly involved cardiovascular activity. These limitations raise the questions whether these findings can be generalized to other environments in which affective states are systematically evoked and whether similar variability is observed with other physiological measures.

To fill this gap, in the current study, we used univariate representational similarity analyses (RSA) to examine the correspondence between multiple physiological responses and subjective affective experiences across different emotion inducing contexts using well-stablished tasks, i.e. passive picture viewing (PPV), passive sound listening (PSL) and mental imagery (I) (Bradley and Lang, 2007). As physiological responses, we focused on four measures previously associated with valence and arousal. Electrodermal (EDA), cardiovascular (heart rate; HR) activity, and startle blink response (in the context of mental imagery; Lang & Bradley, 2007) were used as correlates of arousal, whereas the startle blink response (in the PPV and PSL tasks), the electromyographical (EMG) activity of the *corrugator supercilia*, and HR^1^ were measured as measures of valence (Hamm et al., 1993; Lang et al., 1993; Lang & Bradley, 2007, 2010).

Although univariate analysis may inform about the relation between experienced affect and associated physiological responses, RSA can help to extend these findings by providing information about how changes in these physiological variables correspond to the experienced valence and arousal by comparing the representation similarity matrices (RSMs; pairwise similarity patterns across trials) of changes of the physiological modality (e.g. SCR) to the RSMs of subjective experience (e.g., arousal).

According to the *fingerprint* hypothesis, a correspondence should exist between physiological reactivity and the experienced affect. Therefore, in the three tasks, the RSMs of arousal-related measures (EDA, HR, and startle during I task), should be similar to the RSMs of subjective arousal, whereas RSMs of valence-related correlates (EMG, Startle during PPV and PSL tasks) should correspond to the RSMs of valence ratings.

On the other hand, according to the *populations* hypothesis, the subjective experience of affect in a particular situation should not be related to ongoing physiological demands, but to the similarities to other situations on the abstract properties of affect. Therefore, from this perspective no such correspondence should be expected between physiological reactivity and subjective reports of affect.

## Materials and Methods

### Participants

The sample size was determined using G*Power (Faul et al., 2007, 2009). Given that we had the three tasks, and three categories per task, a total of 57 participants was needed. We decided to collect a few more participants in case recording problems emerged or the quality of the data was compromised. For this reason, a total of sixty-four healthy students (51 women, 13 men; mean age = 23.43) from the University of Potsdam participated in the study in exchange of course credits or financial compensation. All participants had normal or corrected-to-normal vision. Each individual provided written informed consent for a protocol approved by the ethics committee of the University of Potsdam. Prior to the first session, participants were screened and invited to participate if they did not match any of the following exclusion criteria: neurological or mental disorders, brain surgery, undergoing chronic or acute medication, history of migraine and/or epilepsy.

### Material, design and procedure

Participants completed five different study sections. First, participants’ physiological responses were measured during rest for 6 minutes. Afterwards, three affect-inducing tasks were conducted: a passive picture viewing task (PPV), a passive sound listening task (PSL), and an imagery task (I). The order of the three affect-inducing tasks was counterbalanced across participants in six different orders. Thereafter, participants provided subjective valence and arousal ratings of the stimuli used in the three prior tasks, using a computerized version of the self-assessment manikin (SAM; Bradley & Lang, 1994).

For the PPV task, a total of 36 images selected from the *International Affective Picture System* (IAPS; Lang et al., 2008) were presented. Images were preselected based on their normative valence and arousal ratings and grouped in three categories: Pleasant, *mean* (*SD*) valence: 7.00 (0.52), *mean* (*SD*) arousal: 6.74 (0.48); Unpleasant, *mean* (*SD*) valence: 2.61 (0.65), *mean* (*SD*) arousal: 6.6 (0.23); and Neutral, *mean* (*SD*) valence: 5.15 (0.4), *mean* (*SD*) arousal: 3.85 (0.21). The presentation order of the image categories was pseudorandomized, with the restriction that no more than two images of the same category were presented consecutively. The order of the image within each category was fully randomized. Images were presented for 6 seconds with an inter-trial interval (ITI) of 14, 16, or 18 seconds. In half of the trials, an acoustic startle probe (95 dB) white noise was presented 5.5 or 6.5 s after cue onset and 9 s after cue offset in half of the 18-second-long ITIs (total of 6 startles during ITIs). Before the presentation of the stimuli and one minute after the beginning of the task, 2 startle probes were presented without visual foreground to account for startle habituation effects.

For the PSL task, a total of 36 sounds selected from the *International Affective Digital Sounds* (IADS; Bradley & Lang, 1999) were presented. Sounds were preselected based on their normative valence and arousal ratings and grouped in three categories: Pleasant, *mean* (*SD*) valence: 6.96 (0.59), *mean* (*SD*) arousal: 6.84 (0.48); Unpleasant, *mean* (*SD*) valence: 2.99 (0.54), *mean* (*SD*) arousal: 6.7 (0.54); and Neutral, *mean* (*SD*) valence: 5.29 (0.51), *mean* (*SD*) arousal: 4.13 (0.56). The presentation order of the sound categories was pseudorandomized, such that no more than two sounds of the same category were presented consecutively. The order of the sounds within each category was fully randomized. Sounds were presented for 6 seconds with an inter-trial interval (ITI) of 14, 16, or 18 seconds. In half of the trials, an acoustic startle probe (95 dB) white noise was presented 5.5 or 6.5 s after cue onset and 9 s after cue offset in half of the 18-second-long ITIs (total of 6 startles during ITIs). Before the presentation of the stimuli and one minute after the beginning of the task, 2 startle probes were presented without visual foreground to account for startle habituation effects.

For the I task, a total of 18 scripts were selected from different sources: the *Affective Norms for English Text* (ANET; N = 15; Bradley and Lang 2007), from prior studies (Limberg et al., 2011), and from in-house sets were presented. As for images and sounds, scripts were preselected based on their normative valence and arousal ratings (for ANET scripts) in three categories: Pleasant, *mean* (*SD*) valence: 8.28 (0.12), *mean* (*SD*) arousal: 7.48 (0.72); Unpleasant, *mean* (*SD*) valence: 2.21 (0.44), *mean* (*SD*) arousal: 7.69 (0.47); and Neutral, *mean* (*SD*) valence: 5.33 (0.19), *mean* (*SD*) arousal: 3.42 (0.51). The scripts were presented twice, resulting in a total of 36 trials. Participants were instructed to read the scripts during presentation, and after script offset, to keep imagining themselves in the situation as vividly as possible. During the imagery period a circle was presented in the middle of the screen. Before the experimental task started, 3 practice trials were performed to ensure that participants understood the instructions. The presentation order of the script categories was pseudorandomized, again with the restrictions that no more than two scripts of the same category were presented consecutively. The order of the script within each category was pseudorandomized in 8 different orders. Scripts were presented for 12 seconds and followed by a 9-second imagery time window. The inter-trial interval (ITI) was of 14, 16, or 18 seconds. In half of the trials, an acoustic startle probe (95 dB) white noise was presented 5.5 or 6.5 s after onset of the imagery part and 9 s after imagery cue offset in half of the 18-second-long ITIs (total of 6 startles during ITIs). Before the presentation of the stimuli and one minute after task onset, 2 startle probes were presented without visual foreground to account for startle habituation effects.

The subjective valence and arousal ratings were obtained in the lab using the online platform *soscisurvey* (soscisurvey.de). Each stimulus modality was assessed separately in the same order as the previous three experimental tasks (e.g. first images, then imagery script and lastly sounds). Within each stimulus-set, the presentation of the stimuli was randomized. Participants were instructed to indicate on a SAM scale (1-9) how they felt in terms of arousal and valence. For the valence scale, participants were told that lower values (e.g., 3) indicated that they felt unpleasant, unhappy, sad, when perceiving or imagining the stimuli, whereas higher values of the scale (e.g. 6) were to indicate that they felt pleasant, happy, delighted when perceiving or imagining the stimuli. For the arousal scale, participants were instructed to use lower values of the scale if they felt calmed, relaxed, whereas higher number of the scale were to be used if feeling excited, stimulated, aroused when the stimuli were presented or imagined. Following the instructions, two examples were presented for each modality and if no questions arose, the subjective valence and arousal ratings for the previously presented 36 images, 36 sounds or the 18 imagery scripts were assessed.

### Data recording, reduction and response definition

#### Skin Conductance Response

Skin conductance was recorded with the MP-160 BIOPAC system (BIOPAC systems, Goleta, CA) using 2 shielded 8 mm Ag-AgCl sensors applied on the thenar and hypothenar eminence of the palmar surface of the participant’s left hand (Boucsein et al., 2012).

Skin conductance responses (SCRs) were analyzed using Continuous Decomposition Analysis in Ledalab Version 3.4.9 (Benedek & Kaernbach, 2010). The skin conductance data were down sampled to a resolution of 50 Hz and optimized using four sets of initial values. For the PPV and PSL task, the response window was set from 1 to 4 s after stimulus onset. For the I task, the response window was set from 1 to 4 sec after the beginning of the imagery part. Prior to performing the analysis, a visual inspection of the data was carried out to detect possible anomalies during recording.

#### Startle Blink Response

The startle blink response was recorded on the left eye using electromyography (EMG) on the orbicularis oculi muscle. To evoke the startle blink reflex, an acoustic startle probe was presented. The startle probe was a 95 dB, 50 ms burst of white noise with a near instantaneous rise time. The sound was administered in both ears using Audio Technica ATH-PRO700 MK2 headphones. EMG was recorded using two 4mm Ag-AgCl Biopac electrodes filled with electrolyte. Data were collected via MP-160 BIOPAC system (BIOPAC systems, Goleta, CA) at a sampling rate of 2000 Hz. Following EMG startle guidelines (Blumenthal et al., 2005), the data were filtered (high-pass filter of 60 Hz), rectified, and smoothed with a time constant of 10 ms.

Startle blink response were defined using in-house scripts in MATLAB 2020a. All trials were visually inspected and manually scored (c.f., guidelines for startle studies, Blumenthal et al., 2005). Startle responses were considered when starting 20-120 ms after probe onset and peaking within 150 ms. The magnitude of the eye-blink response (in microvolts) was measured from onset to peak. Trials with no detectable blinks were scored as zero responses, and trials with excessive baseline activity or recording artifacts were scored as missing (PPV: 0.8%, PSL: 0.7%, I: 0.5%).

#### Heart rate

Heart rate (HR) was derived from the raw ECG signal recorded continuously during the tasks using a two-lead set-up with the MP-160 BIOPAC system (BIOPAC systems, Goleta, CA). The electrodes were placed on the right arm and left ankle, following the Einthoven’s triangle configuration (lead II). The raw ECG signal was recorded at 2000 Hz. Using in-house scripts, data were visually inspected to detect and correct for artifacts. HR responses were extracted within a window from 2 seconds prior to 6 seconds after stimulus onset in PPV and PSL tasks, and from 2 seconds prior script presentation to 6 seconds after the beginning of the imagery part in the I task. HR responses were averaged in half-second bins (Reyes Del Paso & Vila, 1998) and corrected for the baseline (two seconds prior stimulus presentation), resulting in a total of 13 bins.

#### Corrugator activity

The corrugator EMG activity was recorded using two 4mm Ag-AgCl electrodes filled with electrolyte located over the left corrugator supercilii. Data were collected via MP-160 BIOPAC system (BIOPAC systems, Goleta, CA) at a sampling rate of 2000 Hz. Following recommendations by Fridlund et al, (1984) data were filtered offline (30 Hz high-pass filter) and rectified using a time constant of 20 ms.

7-second segments were extracted from PPV and PSL tasks. The segments included 2 seconds prior to stimulus onset to 6 second after stimulus onset. For the I task, 20-second segments were extracted including 2 seconds prior to stimulus onset to 6 seconds after the beginning of the imagery part. Data were averaged in 100 ms bins, and baseline-corrected using the 2 second prior stimulus onset.

### Data analysis

#### Data exclusion

##### Skin conductance response

Due to anomalies during recording (excessive artifacts or near-to-zero skin conductance levels) 5 participants were excluded from the PPV, 3 from the PSL, and 3 from the I.

##### Startle response

Following the guidelines by Blumenthal et al. (2005), participants with more than 50% of non-responses were excluded from the analysis. This resulted in 3 excluded participants from the PPV, 3 from the PSL, and 2 from the I.

##### Corrugator

Due to technical reasons (lack of EMG module), data from the first eleven participants are missing in the three tasks. Furthermore, due to extreme values, 1 participant was excluded from the PPV, 2 from the PSL, and 3 from the I task.

The analysis of the data was split in two separate sets. The first set of analyses aimed to replicate previous findings on the relation between subjective ratings of valence and arousal and physiological response across participants. In the second set of analyses, RSA was implemented to disentangle between the *fingerprints* and *populations* hypotheses in terms of the relation between valence and arousal and physiological responses. According to the *fingerprints* hypothesis, a relation should exist between the RSMs of physiological reactivity and the arousal- and valence-based RSMs. However, the *populations* hypothesis would predict no such relations.

#### Replication of previous findings

To replicate previous findings, ANOVAs were performed with the within-subjects factors Category (pleasant, unpleasant, and neutral). Post-hoc planned comparisons between categories were conducted using Bonferroni corrections for both subjective ratings and physiological responses.

For EDA, the average phasic driver within the specified response window was derived. To avoid skewed data and extreme values, and following standardized recommendation (Boucsein et al., 2012), the resulting estimates of average phasic driver within the response window were log transformed: log (SCR+1).

For the startle, blink magnitudes were standardized using the 18-startle response presented during trials from a given participant as the reference distribution. The *Z*-standardized responses of each participant were then converted to T scores [50 +(*Z* x10)].

For HR changes, baseline-corrected 500-ms bins were averaged across the 6 second time interval after stimulus onset in PPV and PSL tasks and across 6 seconds time interval after the beginning of the imagery part in the I task.

For the corrugator response, baseline-corrected 100-ms bins were averaged across the 6 second time interval after stimulus onset in PPV and PSL tasks and across 6 seconds time interval after the beginning of the imagery part in the I task.

#### Representational similarity analysis (RSA)

To examine the correspondence between the pattern of physiological response and the subjective experience of valence and arousal, multivariate representational similarity analyses (RSA; Kriegeskorte et al., 2008) were carried out for each physiological variable and task, separately. In RSA, physiological measures evoked by single events are quantified and related to each other. In comparison to univariate methods which may indicate whether averaged events representing one condition differ from another averaged condition, RSA can provide more detailed information about the relation of the physiological reactivity and a particular psychological construct by directly examining the relation between individual responses. One of the most salient characteristics of RSA is its ability to compare physiological reactivity to computational models and behavioral data by comparing representational similarity matrices (RSMs).

Prior studies investigating the physiological reactivity to affective information in the perceptual and imagery domain indicate that SCR and HR changes are related to the degree of experienced arousal (Hamm et al., 1993; Bradley et al., 2001). Thus, RSMs of these physiological measures were compared to RSMs of the subjective experience of arousal. In addition, the startle blink response has also been suggested to capture the subjective experience of arousal during imagery (Bradley & Lang, 2007). For this reason, RSM of the startle blink response in the I task was compared to the RSM of the subject experience of arousal.

Similarly, it has been suggested that the activity of the corrugator in both the perceptual and imagery domain and the modulation of the startle blink response in the perceptual domain are related to the degree of experienced valence (Bradley et al., 2001). Also, some studies have related the HR changes to the experienced valence in perceptual tasks (Brouwer et al., 2013). Thus, RSMs of these physiological measures were compared to RSMs of the subjective experience of valence.

##### Computation of subjective RSMs

Before the subjective RSMs were computed, trials for each participant were sorted based on their own valence or arousal ratings, depending on the physiological response and stimulus domain. Although the stimuli were previously selected to contain instances on the categories of valence (i.e. pleasant, unpleasant, and neutral) and arousal (low, high), we acknowledge that the experienced valence and arousal evoked by a particular event may vary from participant to participant. Thus, rather than determining the sorting of the stimuli based on normative ratings, we used individual subjective ratings, which may be particularly relevant for these psychological constructs (e.g. Jin et al., 2015). As a result, for the tasks tapping on the perceptual domain (PPV and PSL) 36 X 36 subjective RSMs matrices were created in which a value reflecting the similarity (Pearson’s *r*) between two stimuli was contained in each cell. Similarly, 18 X 18 RSMs matrices were created for the I task.

Of note, participants and trials included in the subjective RSMs matched with those included in the physiological RSMs.

##### Computation of the physiological RSMs

Figure 1 depicts the computation of the physiological RSMs using the PPV task and the SCR as example.

**Figure 1.**
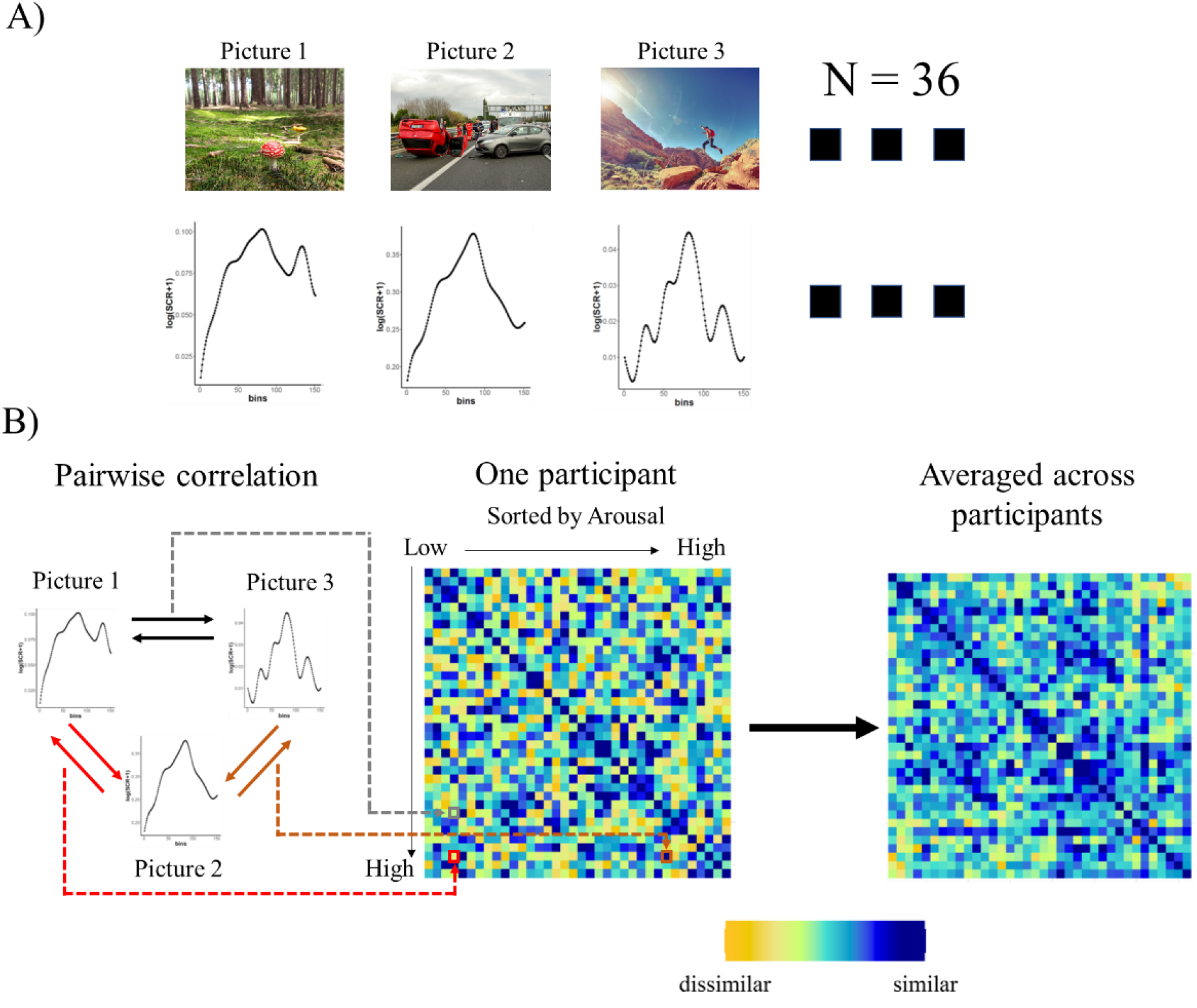
Computation of the physiological RSMs. A) In a first step, the physiological responses were extracted from each trial (36 for the PPV and PSL tasks and 18 for the I task; for the Startle response, 18 startle probes were presented in each task). B) In a second step, each trial was correlated to each other (*Pearson*’s correlation), and sorted in terms of the valence or arousal ratings, resulting in a symmetrical matrix (e.g. 36×36), the representational similarity matrix (RSM), in which the diagonal contained the correlation of each trial with itself and the upper and lower part of it were mirrored. This step was performed for each participant, and then averaged across participants. The averaged RSM was then compared to the subjective RSM, using Spearman’s rank correlations.

#### Skin conductance response

The time-series of the phasic SCR between 1 and 4 seconds after stimulus onset was extracted for each trial and averaged across repeated trials for the I task (150 bins per vector). Next, for each participant, trials were ordered based on subjective arousal ratings and 36 X 36 (i.e. PPV and PSL tasks) and 18 X 18 (I task) RSMs were constructed. Each cell of the RSM consisted of a similarity value (Pearson’s *r*). Thereafter, all RSMs were averaged across individuals for each task.

#### Startle blink response

Because the physiological response of the startle consisted of a single value (t-scored magnitude), no individual RSMs could be computed. Instead, a single RSM across participants was calculated. To do so, firstly each trial was sorted based on the subjective valence ratings for PPV and PSL tasks and based on arousal ratings for the I task for every participant. Thereafter, a 18 X 18 RSM was generated in which each cell reflects the similarity (Pearson’s *r*) between startle magnitudes for a pair of stimuli.

#### Heart rate

A vector of 13 bins (500 ms bins for a total of 6 seconds) was extracted for each trial and averaged across trials for the I task. Next, for each participant, trials were ordered based on subjective arousal ratings, and 36 X 36 (i.e. PPV and PSL tasks) and 18 X 18 (I task) RSMs were constructed. Each cell of the RSM consisted of a similarity value (Pearson’s *r*). Thereafter, all RSMs were averaged across individuals for each task.

#### Corrugator

A vector of 60 bins (100 ms bins for a total of 6 seconds) was extracted for each trial and averaged across trials for the I task. Next, for each participant, trials were ordered based on subjective arousal ratings and 36 X 36 (i.e. PPV and PSL tasks) and 18 X 18 (I task) RSMs were constructed. Each cell of the RSM consisted of a similarity value (Pearson’s *r*). Thereafter, all RSMs were averaged across individuals for each task.

##### Comparing subjective and physiological RSMs

To examine the correspondence between subjective and physiological RSMs Spearman’s rank correlation coefficients were calculated between the physiological RSMs of SCR and HR of the three tasks and the RSM of the startle blink response for the I task and the RSMs of subjective experienced arousal. Similarly, RSMs of the corrugator for the three tasks and of the startle blink response for the PPV and PSL tasks were compared to the RSMs of the subjective experienced of valence. Because the RSMs are mirrored along the diagonal, to avoid inflated correlations, only the values above the diagonal were considered for the correlation. Statistical significance was tested using permutation tests by randomizing the RSM labels (i.e. subjective vs. physiological). The RSM labels were iterated 10,000 times, using Monte Carlo random permutation. For each iteration, the correlation between RSMs was computed. These estimates served as a null distribution against which the actual correlation coefficient was compared. If the actual correlation coefficient was above the 95% of the null distribution, we rejected the null hypothesis that the physiological and subjective RSMs are unrelated.

In addition to the main RSA analysis, we performed two RSM analyses. First, we compared the physiological RSMs to the RSMs based on the complementary affect characteristic (e.g. the SCR-related RSMs, were compared to RSMs in which the relation of each pair of stimuli was based on subjective valence, rather than on subjective arousal). The correlation indexes resulting from both comparisons were then compared. Second, the physiological RSMs of the same variable (i.e. SCR) were compared across tasks (PPV vs. PSL) to investigate whether both tasks were representing similar processes. All analyses were performed with R software.

## Results

### Ratings

Table 1 summarizes the valence and arousal subjective ratings (mean and standard deviation) for each hedonic category and task.

**Table 1.**
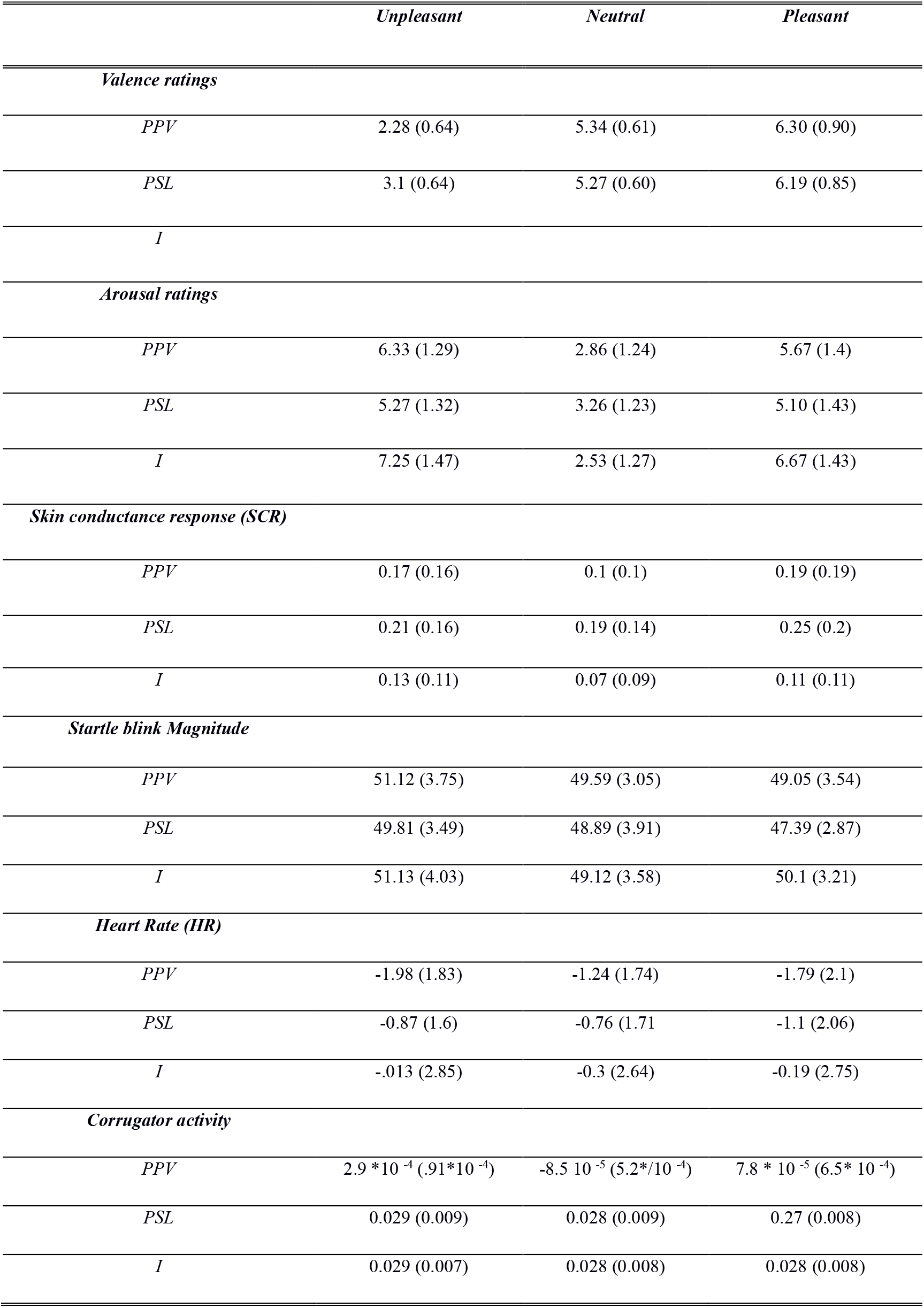
Mean (standard deviation) of subjective valence and arousal ratings and psychophysiological indices for unpleasant, neutral, and pleasant material.

#### PPV Task

For valence ratings, the One-way ANOVA revealed a significant effect of Category, *F* (2,126)= 561.95, *p*<.001, *η_p_*^2^ = 0.89. Bonferroni-corrected, post-hoc comparisons (p = .0167) showed that the valence ratings were lower for unpleasant than neutral images, *t(*63)= −31.53, p <.001, CI[−4.35, −3.78], *d*=3.87, and for neutral compared to pleasant images, *t(*63)= 7.4, p <.001, CI [0.74, 1.27], *d*=.89. Valence ratings for pleasant images were higher than for unpleasant ones, *t(*63)= 28.62, p < .001, CI [3.78, 4.35], *d*= 3.56.

For arousal ratings, the One-way ANOVA revealed a significant effect of Category, *F* (2,126)= 279.0, *p*<.001, *η*_p_^2^ = 0.81. Bonferroni-corrected, post-hoc comparisons (p = .0167) showed that the arousal ratings were higher for both unpleasant *t(*63)= 22.19, p <.001, CI[3.15, 3.78], *d*=2.77, and pleasant pictures compared to neutral ones, *t(*63)= 16.67, p <.001, CI[2.45, 3.11], *d*=2.08. Ratings for pleasant images were lower than for unpleasant ones, *t(*63)= −4.8, p < .001, CI[−0.97, −0.41], *d*= 0.6.

#### PSL Task

For valence ratings, the One-way ANOVA revealed a significant effect of Category, *F* (2,126)= 402.39, *p*<.001, *η*_p_^2^ = 0.86. Bonferroni-corrected, post-hoc comparisons (p = .0167) showed that the valence ratings were lower for unpleasant in comparison to neutral sounds, *t(*63)= −23.48, p <.001, CI[−2.37, −1.99], *d*=2.93. Pleasant sounds were rated more positive than neutral, *t(*63)= 8.31, p <.001, CI [0.69, 1.14], *d*=1.03, and unpleasant sounds, *t(*63)= −23.76, p < .001, CI[2.8, 3.36], *d*= 3.56.

For arousal ratings, the One-way ANOVA revealed a significant effect of Category, *F* (2,126) = 130.28, *p*<.001, *η*_p_^2^ = 0.67. Bonferroni-corrected, post-hoc comparisons (p = .0167) showed that the arousal ratings were higher for both unpleasant, *t(63)=* 14.74, p <.001, CI [1.74, 2.29], *d*=1.84, and pleasant sounds in relation to neutral sounds, *t(63)=* 12.26, p <.001, CI[1.5, 2.01], *d*=1.53, but no differences were observed between pleasant and unpleasant sounds, *t(63)=* −1.77, p = .08, CI[−.48, .03], *d*= 0.22.

#### Imagery Task

For valence ratings, the One-way ANOVA revealed a significant effect of Category, *F*(2,126) = 1,241.72, *p*<.001, *η*_p_^2^ = 0.95. Bonferroni-corrected, post-hoc comparisons (p = .0167) showed that the unpleasant scripts were rated more negatively than their neutral counterparts, *t(*63)= −33.1, p <.001, CI[3.91, −3.47], *d*=4.13, and pleasant scripts were rated more positively than neutral, *t(*63)= 19.41, p <.001, CI [1.99, 2.45], *d*=2.42, and unpleasant ones, *t(*63)= 44.6, p < .001, CI[5.65, 6.18], *d*= 5.6.

For arousal ratings, the One-way ANOVA revealed a significant effect of Category, *F*(2,126) = 308.9, *p*<.001, *η*_p_^2^ = 0.83. Bonferroni-corrected, post-hoc comparisons (p = .0167) showed that both unpleasant, *t(*63)= 20.45, p <.001, CI [4.25, 5.17], *d*=2.44, and pleasant scripts, *t(*63)= 19.55, p <.001, CI[3.7, 4.6], *d*=1.53, were rated more arousing than neutral ones. In addition, pleasant scripts were rated as less arousing than unpleasant scripts, *t(*63)= −3.27, p = .002, CI[−0.92, −0.22], *d*= 0.22.

#### Psychophysiological measures

##### Univariate analysis

Table 1 summarizes the physiological indices (SCR, startle, heart rate and corrugator; mean and standard deviation) for each category and task. Table 2 summarizes the main findings of the univariate analysis.

**Table 2.**
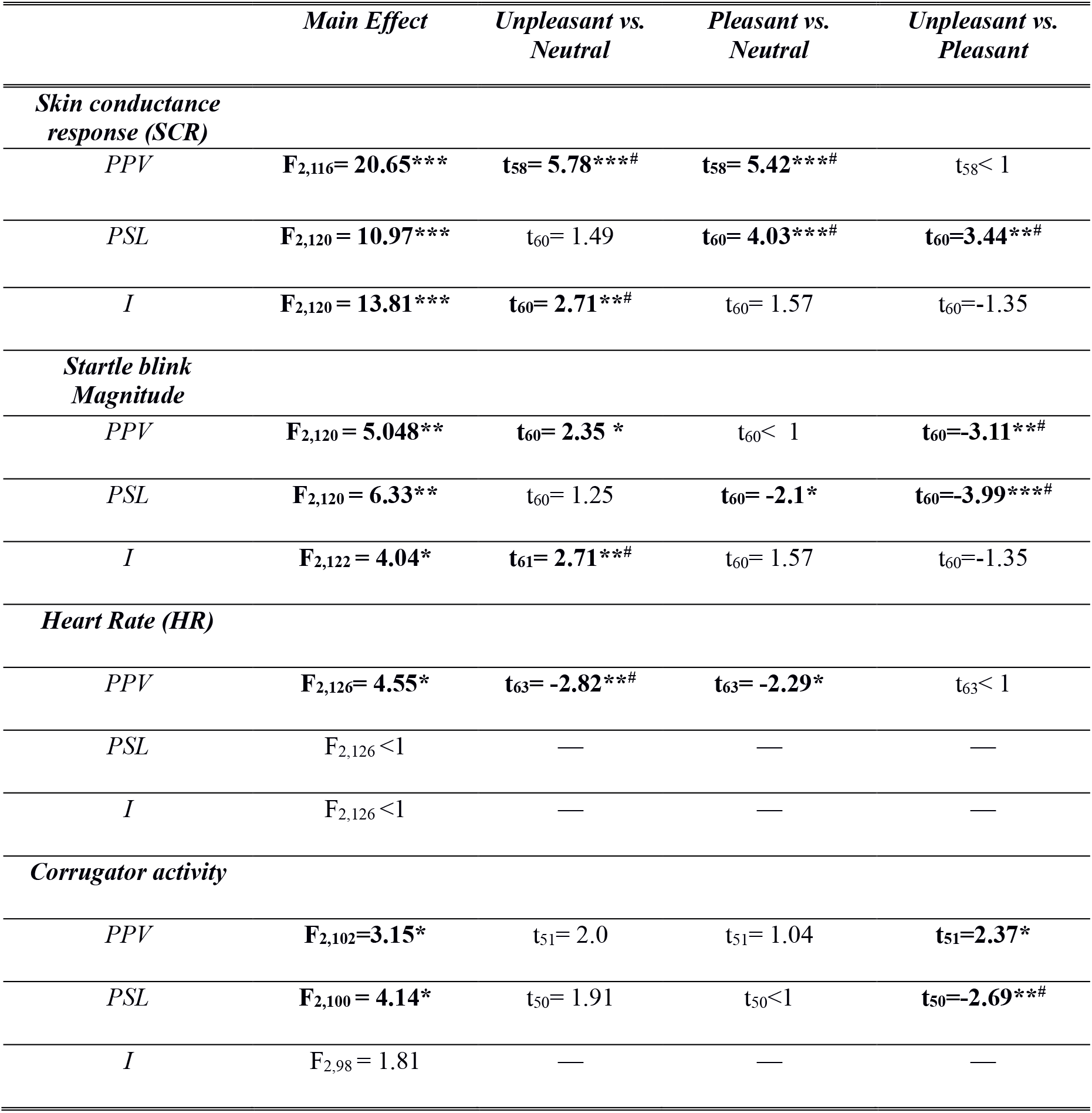
Summary of the main findings of the psychophysiological measures. * p-value < .05; ** p-value <.01, ***p-value <.001, #significant differences after Bonferroni correction (p = .0167). In bold contrasts with p-values lower than 0.05.

###### PPV Task

The One-way ANOVA revealed a significant effect of Category, *F* (2,116) = 20.65, *p*< .001, *η*_p_^2^ = 0.26. Bonferroni-corrected post-hoc comparisons (p = .0167) revealed that both unpleasant, *t(*58)= 5.78, p < .001, CI [.04, .10], *d*=0.75, and pleasant images evoked higher SCRs than neutral ones, *t(*58)= 5.42, p < .001, CI [.05, .13], *d*=0.71, but did not differ between pleasant and unpleasant images, *t*< 1 (Figure 2).

**Figure 2.**
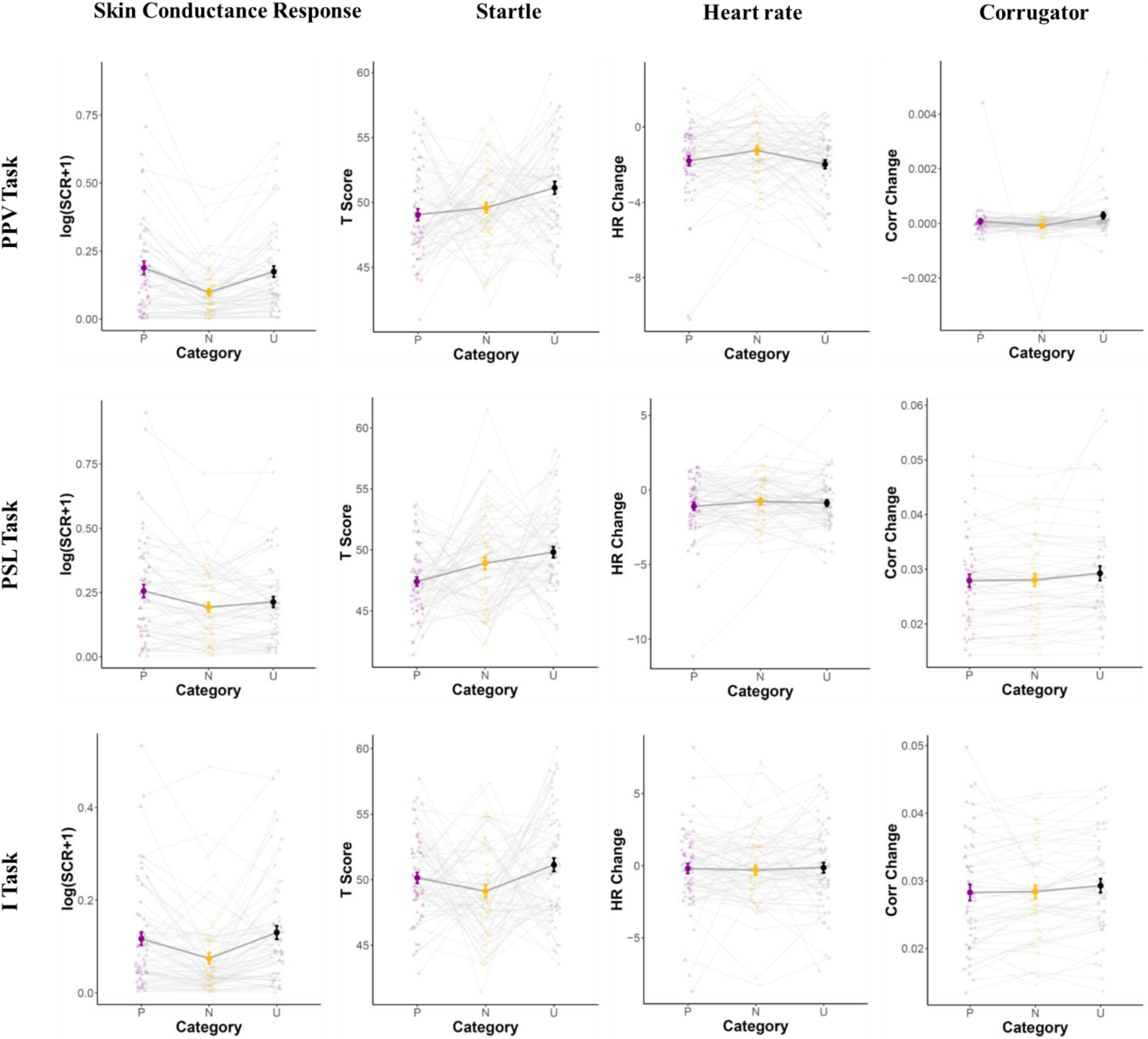
Results of the univariate analysis across category, physiological measure and task. Darker, bigger dots and lines represent the mean response across participants. Error bars represent the standard error. Thinner lighter dots and lines represent responses for each participant. P: pleasant; N: neutral; U: unpleasant. PPV: passive picture viewing; PSL: passive sound listening; I: imagery.

###### PSL Task

For sounds, a significant effect of Category was observed, *F* (2,120) =10.97, *p*< .001, *η*_p_^2^ = 0.15. Bonferroni-corrected post-hoc comparisons (p = .0167) revealed that although the SCR magnitude did not differ between unpleasant and neutral sounds, *t(*60)= 1.49, p = .14, CI [−.012, .05], *d*=0.19, pleasant sounds produced a larger SCR response than neutral, *t(*60)= 4.026, p < .001, CI [.025, .1], *d*=0.52, and unpleasant ones, *t(*60)= 3.45, p = .001, CI [.013, .08], *d*= 0.44 (Figure 2).

###### I Task

For scenes, a significant effect of Category was observed, *F* (2,120)= 13.81, *p*< .001, *η*_p_^2^ = 0.19. Bonferroni-corrected, post-hoc comparisons (p = .0167) revealed that the SCR evoked by both unpleasant, *t(*60)= 4.58, p < .001, CI [.03, .08], *d*=0.58, and pleasant scripts was higher than for neutral scenes, *t(*60)= 3.89, p < .001, CI [.015, .07], *d*=0.49, but did not differ between pleasant and unpleasant scenes, *t(*60)= 3.45, p = .001, CI [.013, .08], *d*= 0.44 (Figure 2).

###### PPV Task

The One-way ANOVA revealed a significant effect of Category, *F* (2,120)= 5.048, *p=* .008, *η*_p_^2^ = 0.08. Bonferroni-corrected, post-hoc comparisons (p = .0167) showed that the startle blink magnitude differed neither between unpleasant and neutral images, *t(*60)= 2.35, p = .028, CI [−.14,3.20], *d*=0.30, nor between pleasant and neutral images, t<1. However, as expected, an inhibition of the startle blink magnitude was observed for pleasant compared to unpleasant images, *t(*60)= −3.11, p = .003, CI [−3.7,-.44], *d*= 0.40 (Figure 2).

###### PSL Task

For sounds, a significant effect of Category was observed, *F* (2,120)= 6.33, *p=* .002, *η*_p_^2^ = 0.10. Bonferroni-corrected, post-hoc comparisons (p = .0167) revealed comparable startle blink magnitudes between unpleasant and neutral sounds, *t(*60)= 1.25, p = .21, CI [−.87, 2.70], *d*=0.16, and between pleasant and neutral sounds, *t(*60)= −2.1, p = .04, CI [−3.25, .24], *d*=0.27. However, the startle blink magnitude was again significantly reduced for pleasant compared to unpleasant sounds, *t(*60)= −3.99, p < .001, CI [−3.9,-.92], *d*= 0.51 (Figure 2).

###### I Task

For imagery scenes, a significant effect of Category was observed, *F* (2,122)= 4.04, *p=* .019, *η*_p_^2^ = 0. 06. Bonferroni-corrected, post-hoc comparisons (p = .0167) revealed that the startle blink magnitude was potentiated for unpleasant compared to neutral scenes, *t(*61)= 2.71, p = .009, CI [.19, 3.84], *d*=0.34, but no differences were observed between pleasant and neutral scenes, *t(*61)= 1.57, p = .12, CI [−.56, 2.61], *d*=0.2, or between pleasant and unpleasant scenes, *t(*61)= −1.35, p = .17, CI [−2.7, .8], *d*= 0.17 (Figure 2).

###### PPV Task

The One-way ANOVA revealed a significant effect of Category, *F* (2, 126)= 4.55, *p=* .012, *η*_p_^2^ = 0.07. Bonferroni-corrected, post-hoc comparisons (p = .0167) revealed that the HR was reduced during unpleasant compared to neutral images, *t(*63)= 2.82, p = .006, CI [−1.37, −.095], *d*=0.35, but no differences were found between pleasant and neutral images, *t(*63)= −2.29, p = .025, CI [−1.13, .04], *d*=0.28, or between unpleasant and pleasant images, *t*< 1 (Figure 2).

###### PSL Task

The One-way ANOVA did not reveal a significant effect of Category, *F*<1 (Figure 2).

###### I Task

For scenes, the One-way ANOVA did not reveal a significant effect of Category, *F*<1 (Figure 2).

###### PPV Task

The One-way ANOVA revealed a significant effect of Category, *F* (2, 102)= 3.15, *p=* .047, *η*_p_^2^ = 0.06. Bonferroni-corrected post-hoc comparisons (p = .0167) revealed that the Corrugator activity did not differ between unpleasant and neutral images, *t(*51)= 2.0, p = .05, CI [−8.61×10^−5^, 8.33 x10^−4^], *d*=0.27, pleasant and neutral images, *t(*51)= 1.04, p = .3, CI [−2.2x x10^−4^, 5.4 x10^−4^], *d*=0.14, or unpleasant and pleasant ones, *t(*51)= 2.37, p = .021, CI [−4.3x x10^−4^, 8.05 x10^−4^], *d*=0.32 (Figure 2).

###### PSL Task

The One-way ANOVA revealed a significant effect of Category, *F* (2, 100)= 4.14, *p=* .019, *η*_p_^2^ = 0.08. Bonferroni-corrected post-hoc comparisons (p = .0167) revealed that the corrugator activity did not differ between unpleasant and neutral sounds, *t(*50)= 1.91, p = .06, CI [−3.5×10^−4^, 2.82 x10^−3^], *d*=0.27, or pleasant and neutral sounds, *t*<1. However, the corrugator activity was decreased for pleasant compared to unpleasant sounds *t(*50)= −2.69, p = .009, CI [−2.6x x10^−3^, −1.1 x10^−4^], *d*=0.37 (Figure 2).

###### I Task

For scenes, the One-way ANOVA did not reveal a significant effect of Category, *F* (2, 98)= 1.81, *p=* .17, *η*_p_^2^ = 0.04 (Figure 2).

In summary, emotional category modulated the SCR and startle blink response in all of the three tasks. An emotional modulation of the HR was, however, only observed in the PPV. Similarly, a significant modulation of corrugator activity was particularly present in the perceptual tasks. These results are mostly in line with prior studies (Bradley et al., 2001; Lang et al., 1999; Hamm et al., 1993 Vrana & Rollock, 2002).

##### Representational Similarity Analysis

###### Skin conductance

To test for the similarities between the subjective experience of arousal and the SCR, the subjective arousal RSMs and the SCR RSMs were compared to each other for each task, separately. For the PPV task, random permutation tests showed a significant correlation between the subjective experience of arousal and the SCR, r= .171, p =.003. However, no significant correlations were found for the PSL and the I tasks (PSL: r= .039, p = .27; I: r = .029, p = .51; Figure 3).

**Figure 3.**
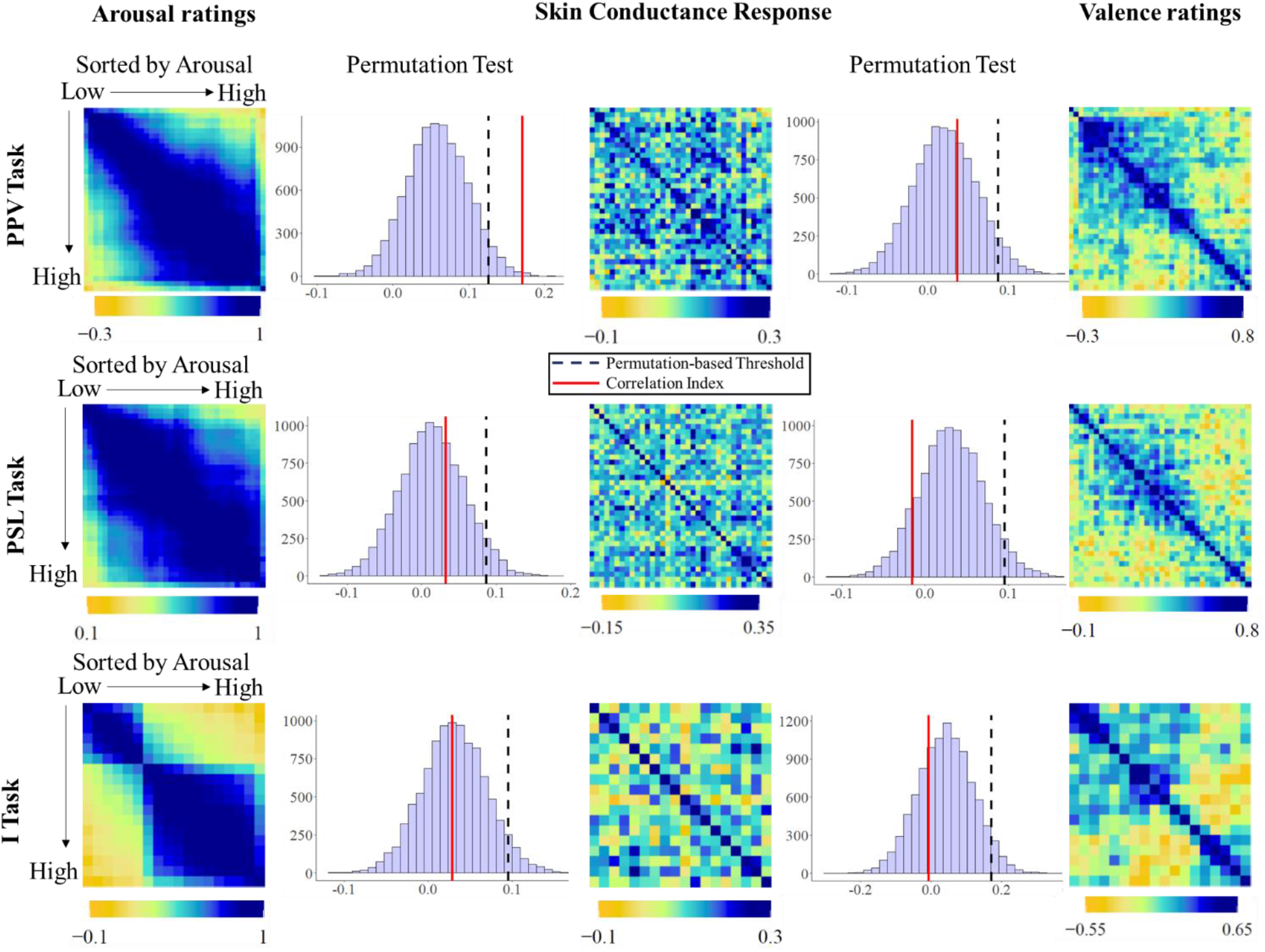
Results from the RSA of SCR. The arousal-sorted RSMs of subjective arousal (right) and valence (left) were compared to RSM of SCR (middle). For the PPV, a significant relation was observed between the arousal-based RSM and the RSM of SCR, as indicated by the random permutation tests. No such relation was observed for the PSL or I task. Similarly, no significant relation was observed between the valence-based RSMs and the RSMs of SCR in any of the tasks.

To test the specificity of the relation between SCR RSMs and those of arousal ratings, the SCR RSMs were also compared to RSMs based on valence ratings. Random permutation tests showed no significant relationship between SCR RSMs and valence RSMs (PPV: r=.036, p =.35; PSL: r= −.015, p =.88; I: r = −.008, p = .73). Indeed, for the PPV, SCR RSMs were more strongly related to the arousal RSMs than to the Valence RSMs (z = 3.17, p = 0.001), but not for the PSL (z = 0.37, p = 0.71) or the I tasks (z = 0.12, p = 0.9).

To test for the similarities between SCR RSMs from different tasks, we also compared the SCR RSMs for the PPV and PSL, which revealed no significant relation (r= .02, p = .73).

###### Startle Blink Reflex

To test for the similarities between the subjective experience of valence and the startle blink reflex, the subjective valence RSMs and the startle RSMs were compared for the PPV and the PSL task, separately. Random permutation tests did not reveal any significant correlation (PPV: r = −0.03, p =.85; PSL: r= −.03, p =.58; Figure 4).

**Figure 4.**
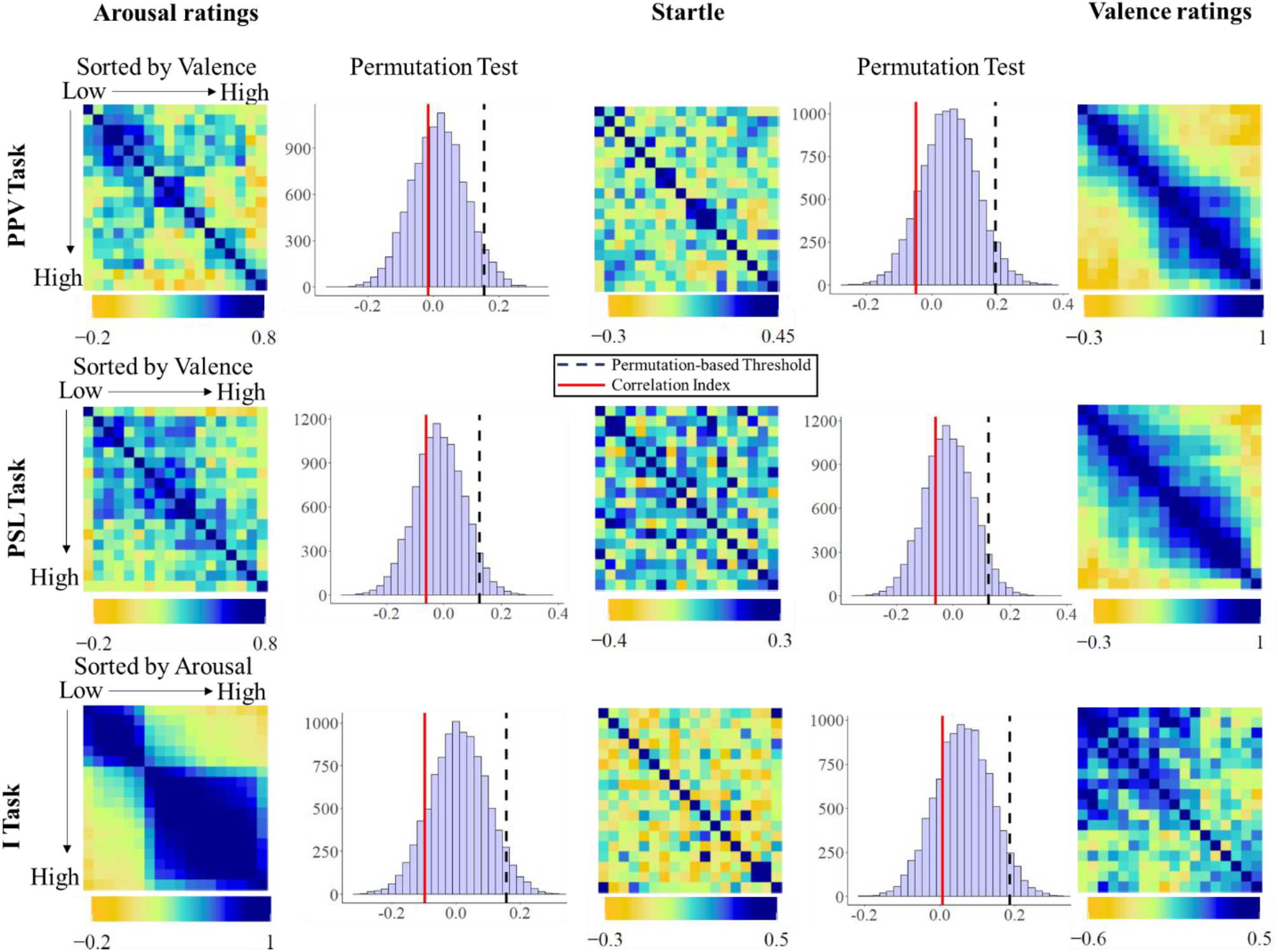
Results from the RSA of the startle blink reflex. RSMs of subjective arousal (right) and valence (left) were compared to RSM of startle (middle). For the PPV and PSL tasks, the trials were sorted by valence, whereas for the I task, the trials were sorted by arousal. No significant relation was observed for any of the comparisons in any of the tasks.

Because previous studies revealed an arousal-mediated modulation of the startle reflex during mental imagery (Bradley & Lang, 2007), the startle RSMs during the I tasks were sorted by arousal (instead of by valence as for in the PPV and PSL tasks) and were compared to the RSMs of the subjective ratings of arousal, which, however, showed no significant relation (r= −.11, p = .89);

The startle RSMs were also compared to RSMs based on complementary affect characteristics. Random permutation tests showed no significant relationship between startle and arousal RSMs for the PPV (r= −.03, p =.66) and PSL tasks (r = −.05, p =.63), and between startle and RSMs for the I task (r= −.02 p= 0.85). No differences were observed between the startle RSMs and the RSMs of valence and arousal (PPV: z = −0.2, p = 0.84; PSL: z =.39, p = .69; I: z= −.58, p = .56).

To test for the similarities between startle RSMs from different tasks, the startle RSMs for the PPV and PSL were also compared, which, however, showed no significant relation (r= −.03, p = .64).

###### Heart rate

To test for the similarities between the subjective experience of arousal and HR, the HR RSMs (sorted by arousal) and the subjective arousal RSMs were compared for each task, separately. Random permutation tests showed no significant relationship between HR and arousal RSMs (PPV: r=.08, p =.22; PSL: r= −.01, p =.58; I: r= .13, p =.25). The HR RSMs were also compared to arousal-sorted RSMs based on subjective valence. Random permutation tests showed no significant relationship between HR and valence RSMs (PPV: r=.05, p= .25; PSL: r= 0.0, p = .66; I: r =-.02, p =.83). No differences were observed for the relation between the Startle RSMs and the RSMs for valence and arousal (PPV: z = −0.52, p = 0.59; PSL: z =.01, p = .83; I: z= 1.09, p = .28; Figure 5).

**Figure 5.**
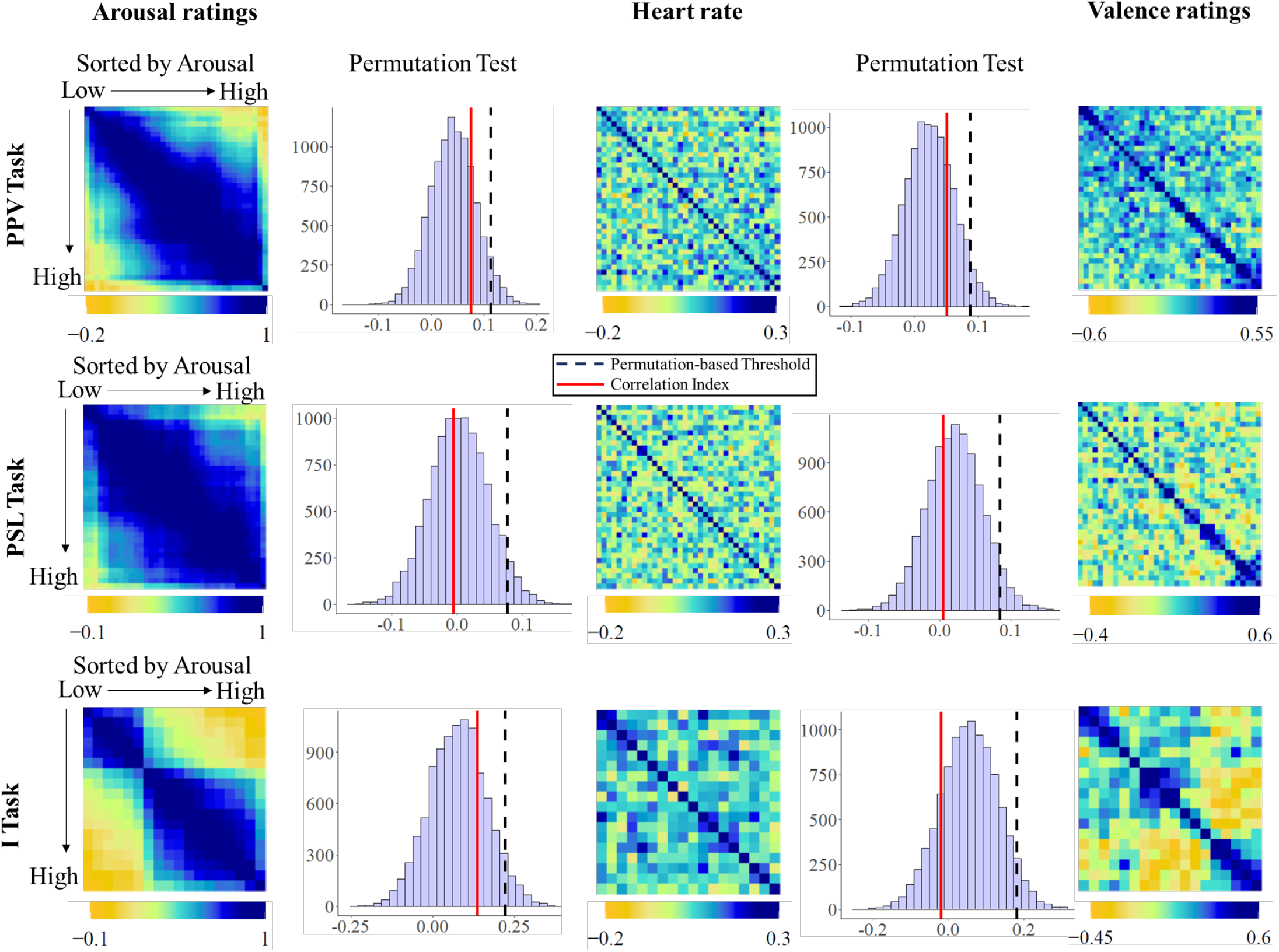
Results from the RSA of heart rate. The arousal-sorted RSMs of subjective arousal (right) and valence (left) were compared to RSM of HR (middle). No significant relation was observed for any of the comparisons in any of the tasks.

Additionally, to test for the similarities between the subjective experience of valence and HR, the HR RSMs (sorted by valence) and the subjective valence RSMs were compared for each task, separately. Random permutation tests showed no significant relationship between HR and arousal RSMs (PPV: r= −.069, p =.99; PSL: r= −.02, p =.92; I: r =.042, p = .93).

The HR RSMs were also compared to valence-sorted RSMs based on subjective arousal. Random permutation tests showed no significant relationship between HR and valence RSMs (PPV: r=-.02, p =.87.; PSL: r=-.04, p =0.76; I: r =-.03, p =.61). No differences were observed for the relation between the HR RSMs and the RSMs for valence and arousal (PPV: z = −1.3, p = 0.19; PSL: z =0.15, p = .87; I: z=1.02, p = .31).

To test for the similarities between HR RSMs from different tasks, we also compared the HR RSMs sorted by arousal and by valence for the PPV and PSL but found no significant relation (arousal: r= .07, p = .08; valence: r= −.02, p = .81).

###### Corrugator

To test for the similarities between the subjective experience of valence and corrugator activity, the corrugator RSMs and the subjective valence RSMs were compared for the three tasks, separately. Random permutation tests did not revealed any significant correlation in any of the tasks (PPV: r = 0.05, p = 0.72; PSL: r= .04, p =.43;I = r= .00, p = .98; Figure 6).

**Figure 6.**
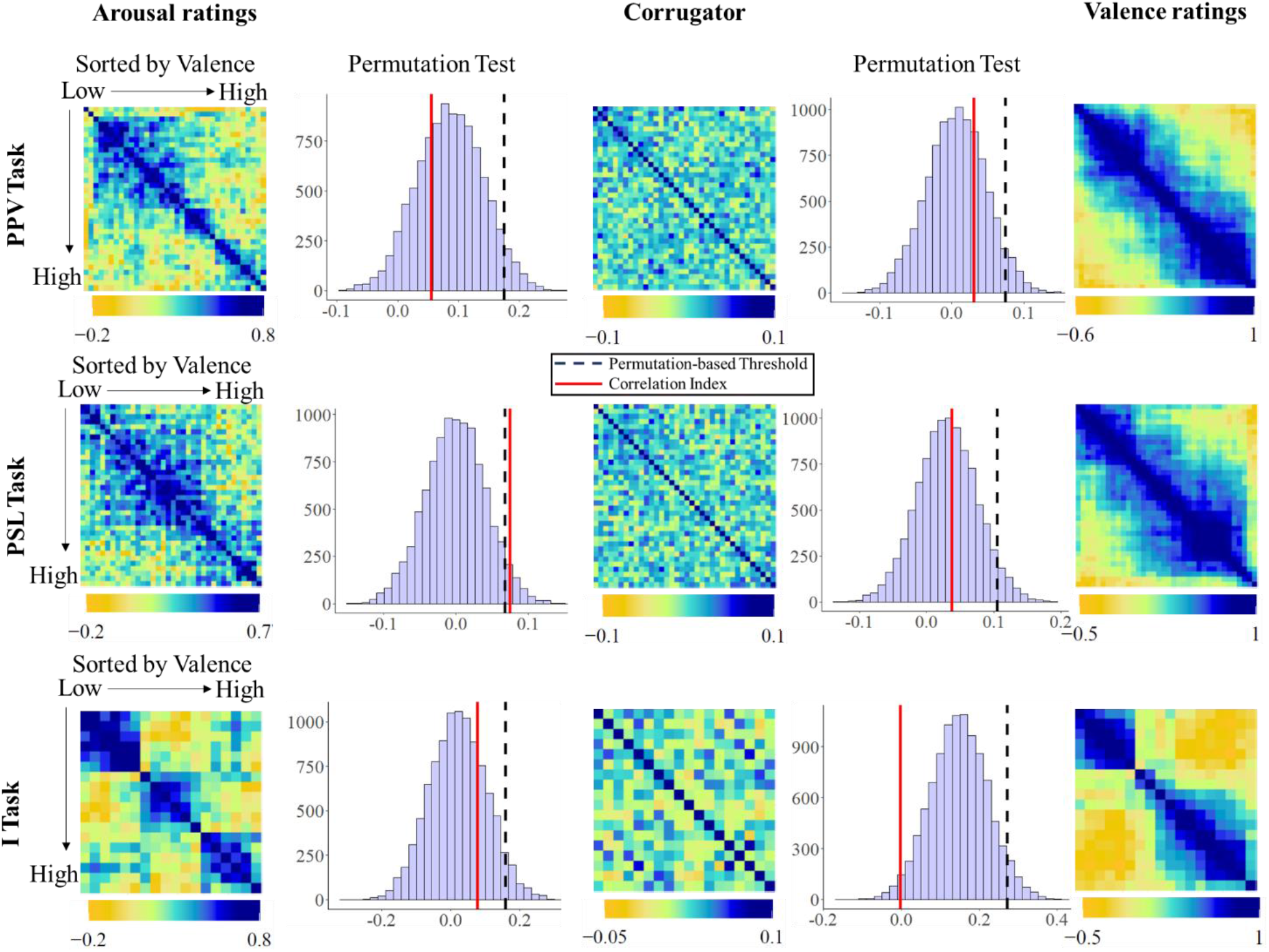
Results from the RSA of corrugator activity. The valence-sorted RSMs of subjective arousal (right) and valence (left) were compared to RSM of corrugator activity (middle). Whereas no significant relation was observed between the valence-based and corrugator RSMs for any of the tasks, a significant relation between the arousal-based RSM and the RSM of corrugator activity was observed for the PSL task.

The corrugator RSMs were also compared to valence-sorted RSMs based on arousal ratings. Random permutation tests showed no significant relationship between corrugator and arousal RSMs for the PPV (r=0.03, p =.28), the PSL (r=.07, p =.03), and the I task (r= .08, p=.25). No differences were observed in the relation between the Corrugator RSMs and the RSMs for valence and arousal (PPV: z = 1, p = 0.31; PSL: z =-1.3, p = .25; I: z= −1.1, p = .27).

Also, to test for the similarities between corrugator RSMs from different tasks, we compared the PPV and PSL tasks, but no significant relation was found (r= .01, p = .32).

In summary, a significant correspondence between the subjective affective experience and physiological measures only emerged for the SCR and arousal in the PPV task. However, no such relationship was observed for the PSL or I task. Also, no significant relationship was observed for the others physiological variables indicating that, except for one case (SCR and arousal during picture viewing), no similarity was found between the pattern of physiological responses and the subjective measures of affect between individuals.

## Discussion

The current study aimed at providing new insights into the debate about whether specific subjective affective states are associated with a particular physiological response. In three different affect-inducing tasks, we applied RSA to investigate the trial-by-trial similarities between subjective affect (arousal and valence ratings) and physiological reactions. RSA allows to determine whether the output of two different modalities is influenced by the same process, by comparing their RSMs. If a link between subjective affect and physiological reactivity exists, as proposed by the *fingerprint hypothesis*, the subjective and physiological RSMs should be similar to each other. However, if the physiological responses are independent of the experienced affect, as stated by the *populations hypothesis*, no similarities should emerge between the subjective and physiological RSMs. We observed that the RSM of SCR was significantly related to the arousal-based RSM in the PPV task. However, none of the RSMs of the other physiological measures showed a significant relation with the RSM of the valence and arousal ratings in any of the tasks. These findings therefore tend to support the *populations hypothesis*, suggesting that there is no clear match between the evoked physiological responses and the experienced subjective affect.

The present results are in line with recent findings indicating that the physiological changes evoked in the context of experienced basic emotions are heterogenous and variant rather than specific and consistent (Siegel et al., 2018; Hoemann et al., 2020). In a recent meta-analysis of 202 studies, Siegel and colleagues (2018) investigated whether “basic” emotions (e.g. happiness anger, fear, surprise, disgust, sadness) are associated with specific physiological responses that can distinguish them from each other. The authors observed a moderate amount of variability between the physiological changes evoked by each emotion across studies which resulted in heterogeneous patterns of physiological responses that could not be clearly distinguished across emotions, supporting the *populations hypothesis*. Our results extend these findings from basic emotions to the valence and arousal properties of affect, demonstrating variability in the physiological responses evoked by events that produced a similar affective experience. An exception was the case of the SCR in relation to arousal in the PPV. However, the fact that this relation was not observed in the PPV and I tasks and that the RSM of SCR was not similar between the PPV and PSL tasks, suggests that the SCR-arousal relation might be dependent on the visual characteristics of the task. One key difference between the pictorial and the acoustic stimuli is that whereas the images remain constant throughout their presentation in the screen, the sounds changed dynamically (Bradley and Lang, 2000), which may have produced a stronger variation in the physiological reactivity. The question arises whether the specificity of the relationship for pictorial stimuli is exclusively related to the evoked arousal experience, due to other uncontrolled characteristics of the stimuli (i.e. static vs. dynamic nature) or due to a combination of both. To determine the specificity of the SCR-arousal relationship to the visual domain future studies focused on replicating the current findings that also control for other potential influential attributes could help to better delineate the relation between subjective arousal and the SCR above and beyond other confounds.

Given that the *population hypothesis* conceptualizes the brain as an entity that acts based on active inference (Friston, 2010; Seth, 2013; Smith et al., 2019) like the theory of constructed emotions (Barrett, 2017; Barrett & Simmons, 2015), the current findings could be better understood within this framework. From this perspective, the final goal of the brain is guaranteeing equilibrium or homeostasis in the body by means of regulating the energy needs. To perform an optimal regulation, the brain builds internal models of the surrounding world and the inner body. These models are constructed and refined based on the regularities acquired through past experiences. Using these models, the brain unveils predictions of what the energy costs will be in the upcoming situations (e.g. going to the beach) and activates a cascade of viscero-motor changes to organize the resources based on these predictions (increase of the heartbeat, and sweating). This compound of changes is then stored as part of hierarchical, multi-dimensional abstractions (e.g. “a pleasant and exciting event”). With time, new lived experiences that evoked different physiological responses (e.g. decrease of heartbeat when anticipating a sport event) are incorporated to these abstractions, increasing the range of physical and environmental features encompassed in them, and strengthening the interconnectivity to other similar abstractions. Due to that, the physiological responses evoked by different instances of the same abstraction (e.g. pleasantness) can be notably different, and similar physiological responses (increase of heart rate) can be related to two different abstractions (pleasantness and unpleasantness). As a result, the categories of affect (pleasantness, unpleasantness, calming, exciting) are composed by instances with shared abstract regularities (beach, sport event) that do not necessarily match in terms of their physical and contextual characteristics. As a consequence, the resources needed for coping with each of these instances may be remarkably different, resulting in different physiological responses.

Several aspects need to be considered when interpreting the present results. The evaluation of affect in terms of valence and arousal may fluctuate from moment to moment and depend on the previously encoded material (Asutay et al., 2019), as well as on whether the stimulus is novel (i.e. seen for the first time) or familiar (i.e. seen the second time; Baskin-Sommers et al., 2013). In the current design, the physiological reactions and subjective ratings were recorded at different time points. It could, therefore, be that experienced affect differed during the emotion induction tasks and the ratings. It is also important to note that participants performed the three tasks in the same session, one by one. Although the order of the task was counterbalanced across participants, it could be that the physiological responses to the evoking material changed throughout the experimental session (e.g. due to habituation, fatigue, prediction etc.). We, however, consider these methodological aspects as minor given the replication of previous findings using univariate analysis (Bradley et al., 2001; Bradley and Lang, 2000; Bradley & Lang, 2007), but we acknowledge that this could have an influence on the correspondence between physiological reactivity and ratings responses.

In summary, in the present study we used RSA to examine the correspondence between subjective affective experience, as measured by valence and arousal ratings, and the associated physiological reactivity in three affect-inducing tasks. Most of the physiological responses across the three tasks showed no correspondence with the expected subjective experience, suggesting that the evoked physiological changes vary across similar experienced instances of affect (as stated by the *populations hypothesis*). These findings do not underestimate the importance of physiological changes in the experience of affect but invite to reframe the relation between physiology and affect from invariant and homogeneous to variant and context-dependent (Hoemann et al., 2020). New perspectives adopting an active inference approach (Barrett & Satpute, 2019; Barrett & Simmons, 2015; Friston, 2010; Seth, 2013) seem promising to help understanding how physiological reactions are implemented on the internal models to create and/or update the existing abstractions conceptualizing affect.

## Research Disclosure Statements

- All dependent variables or measures that were analyzed for this article’s target research question have been reported in the Methods section(s)
- All levels of all independent variables or all predictors or manipulations, whether successful or failed, have been reported in the Method section(s)
- The total number of excluded observations and the reasons for making those exclusions (if any) have been reported in the Method section(s)

## Footnotes

1 Because previous studies have shown an arousal- and valence-mediated modulation of HR changes (Bradley et al., 2001; Brouwer et al. 2013), we investigated the relation of HR changes with both valence and arousal ratings.

